# UltraSEQ: a universal bioinformatic platform for information-based clinical metagenomics and beyond

**DOI:** 10.1101/2022.08.24.505213

**Authors:** Bryan T Gemler, Chiranjit Mukherjee, Carrie Howland, Patrick A Fullerton, Rachel R Spurbeck, Lindsay A Catlin, Anthony Smith, Angela T Minard-Smith, Craig Bartling

**Author notes:** Bryan T Gemler and Chiranjit Mukherjee contributed equally to the manuscript. Author order was determined alphabetically. Craig Bartling is the corresponding author.

## Abstract

Applied metagenomics is a powerful emerging capability enabling untargeted detection of pathogens, and its application in clinical diagnostics promises to alleviate the limitations of current targeted assays. While metagenomics offers a hypothesis-free approach to identify any pathogen, including unculturable and potentially novel pathogens, its application in clinical diagnostics has so far been limited by workflow-specific requirements, computational constraints, and lengthy expert review requirements. To address these challenges, we developed UltraSEQ, a first-of its kind metagenomics-based clinical diagnostics and biosurveillance tool that is accurate and scalable.

Here we present results for evaluation of our novel UltraSEQ pipeline using an *in silico* synthesized metagenome, mock microbial community datasets, and publicly available clinical datasets from samples of different infection types, and both short-read and long-read sequencing data. Our results show that UltraSEQ successfully detected all expected species across the tree of life in the *in silico* sample and detected all 10 bacterial and fungal species in the mock microbial community dataset. For clinical datasets, even without requiring dataset-specific configuration settings changes, background sample subtraction, or prior sample information, UltraSEQ achieved an overall accuracy of 91%. Further, we demonstrated UltraSEQ’s ability to provide accurate antibiotic resistance and virulence factor genotypes that are consistent with phenotypic results.

Taken together, the above results demonstrates that the UltraSEQ platform offers a transformative approach to microbial and metagenomic sample characterization, employing a biologically informed detection logic, deep metadata, and a flexible system architecture for classification and characterization of taxonomic origin, gene function, and user-defined functions, including disease-causing infection.

## Introduction

Traditional clinical microbiology based diagnostic tests rely on targeted methods that can only detect one to a few preselected organisms through the use molecular methods (e.g., quantitative polymerase chain reaction (qPCR) or antigen-detection methods) or slow, culture-based methods. Although widely used today, these methods have several limitations, especially for infections caused by more than one etiological agent, novel pathogens, and unculturable organisms. Further, these tests are often single-plex, requiring multiple tests to be run prior to diagnosis. Due to these limitations, the rate of unknown etiology of infection has been reported to be >50% for diseases such as pneumonia and encephalitis [1, 2]. This rate of unknown etiology is corroborated by our initial analysis of commercial health claims of 2021 International Classification of Disease (ICD) codes, which suggests that despite >$5 billion charged in 2021, as high as 80% of pneumonia cases (average of inpatient and outpatient) were coded with an ‘unspecified organism’ ICD code [3].

Massive development in sequencing technologies have made it possible to apply metagenomic methods to clinical diagnostics, which may alleviate the issues described above. Metagenomics offers a hypothesis-free approach to identify any pathogen, including unculturable and potentially novel pathogens in a massively-multiplexed assay that requires little to no wet lab assay development for new etiological agents. The approach is largely unbiased, can work well with very low biomass samples, can often replace invasive sampling regiments, and ultimately lead to better patient outcome and reduce antibiotics misuse through quick, accurate determination of the disease cause (refs). Clinical metagenomics is rapidly moving from research to the clinic, with commercial offerings for diseases such as sepsis, respiratory disease, and meningitis/encephalitis [4]. However, current offerings are limited to a specific disease type, sequencer workflow, and/or require laboratory-specific controls, thus limiting their widespread adoption.

The limitations associated with current clinical metagenomic offering result from the fact that the backend bioinformatic pipelines are optimized for the specific parameters described above, resulting in an excess of unmaintained, redundant, and niche tools that lack standardization and explainable output. For pathogen identification, several different classifiers exist, including those that leverage alignment-based (both nucleic acid and protein) and k-mer-based approaches [5, 6]. Classification can be provided on the individual sequence level (binning) as well as the whole dataset level (profiling). While k-mer based methods are fast, efficiency must be weighed against optimization of k-mer sizes with each database update as well as the lack of granularity in predictions due to exact matching (i.e., important variable sequence information relevant to clinical settings may be lost [7]). Thus, k-mer based approaches can be highly specific, but suffer from low sensitivity, particularly for identification of novel pathogens such as viruses with high mutation rates [8]. Most clinical metagenomic tools have adopted rapid alignment routines to achieve high sensitivity. However, such routines often result in over identification of organisms in samples, many of which may be false positives. Tools may overcome these high false positive rates by using background subtraction methods and thresholding on various settings that are optimized for each specific workflow. Often conservative predictions are made at higher taxonomic levels (e.g., genus-level) that are less informative for clinical applications.

To address all of these challenges, we developed UltraSEQ, a first-of its kind metagenomics based clinical diagnostic tool that is accurate, scalable, and provides an information-based approach for clinical research applications. As described above, many pathogen identification bioinformatic routines suffer from high false positive rates, the inability to identify organisms not included in their reference database, and either complexity associated with interpretation of results or non-explainable ‘black-box’ answers. In contrast, UltraSEQ uses a novel, information-based approach that leverages a fast aligner that can handle both DNA and protein database to make sample-level predictions (including taxonomic profiling) at the most specific taxonomic levels possible given the information in the sample and the database(s) used. UltraSEQ was built from the ground up to make predictions for regions of sequences (including taxonomic binning), full sequences, and collections of sequences (i.e., a sample) without complicated user settings and the necessity for background subtraction. This novel approach enables accurate, evidence-based predictions for diagnosis as well as functional characterization of a sample, including virulence factor and antibiotic resistance profiles. Predictions are backed by our curated database that provides end users with additional contextual metadata of predictions, such as whether the identified pathogen typically or rarely causes disease, is a potential normal flora contaminant, as well as other user-defined characteristics. In our previous work, we described the development of this database [9] that uses a function-centric approach to distinguish between pathogenic and non-pathogenic organisms with increased confidence over the state-of-the-art. Here we expand upon this approach to leverage this database by building the flexible UltraSEQ bioinformatic platform and demonstrate its utility for clinical metagenomic applications that go *beyond simple taxonomic prediction*. We demonstrate UltraSEQ’s ability to handle a variety *of different sample types, sequencing platforms, and laboratory workflows and UltraSEQ’s superior performance compared to several other bioinformatic platforms.*

## Methods

### Overall UltraSEQ Architecture and Information Flow

The overall UltraSEQ pipeline is illustrated in **Figure 1**. UltraSEQ handles various input file types and relies on high quality sequence regions for downstream predictions. Each sequence has the option to go through all or some of the services described below, including a quality assurance pre-processing service, an aligner service, a query mapper service, context services, and prediction services. Collections of sequences can then undergo additional predictions including taxonomic composition prediction (metagenomics module), rules engine(s), and reporting services. A high-level summary of the services used in this study are located below, with details of those services used in this study provided in **Supplemental Material A: Detailed UltraSEQ Services and** other services not used during this study described in **Supplemental Material B: Other UltraSEQ Services** (for completeness sake). UltraSEQ has a modular architecture with non-restrictive deployment, including both local, high performance computing cluster, and cloud-based deployment. UltraSEQ is currently deployed on Amazon Web Services (AWS) in a secure, scalable computing environment complete with graphical user interface to submit samples and download result reports and is available for use through contacting the corresponding author.

**Figure 1.**
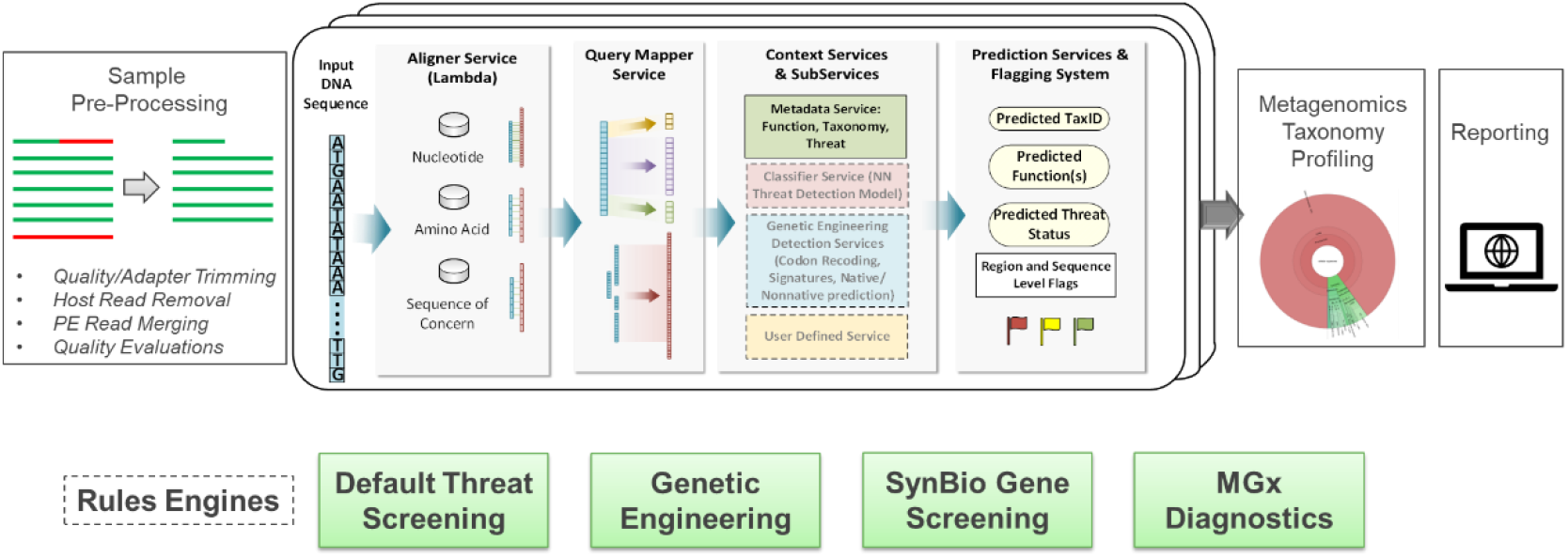
UltraSEQ Pipeline graphical representation.

#### Preprocessing Service

UltraSEQ’s preprocessing routine includes steps to trim low quality sequence regions, remove adapter sequences, remove duplicate sequences, (optionally) merge paired end reads, and (optionally) remove host sequences [10–16]. Different routines were applied to short read (Illumina and IonTorrent) datasets and long read (Nanopore) datasets as detailed in **Supplemental Material A**. Further, to reduce cloud compute costs, enhance run-times, and provide better comparisons across datasets, subsampling was optionally performed prior to the alignment service as described in **Supplemental Material A**.

#### Aligner Service

Each sequence is rapidly aligned using LAMBDA2 version 1.9.5 [17] against selected databases, although other aligners can be used if desired. LAMBDA2 enables alignment against both protein and nucleotide databases, but the results for this study leveraged only protein databases, including the Uniref100 protein database (built April 2021) and Battelle’s Sequence of Concern (SoC) protein database [9].

#### Query Mapper Service

This service maps regions within query sequences (i.e., portions of the query sequence) to identify high quality alignment regions as well as chimeric reads / out-of-context DNA sequences. For each query sequence, the positions of alignment starts and stops from high quality alignments were compiled. A K-means clustering approach was used to identify positions in the query sequence with a high abundance of alignment starts/stops, which are subsequently used to identify regions of the query sequence.

#### Context Services and Subservices

These services generate contextual information and passes information to downstream services. Information from the context services is passed to the Prediction Services and Flagging System (Rules Engine) as described below.

#### Region-based Prediction Subservices

For each region identified from the Query Mapper Service, UltraSEQ predicts the taxonomy, function (gene ontology terms), and threat associated with the region. For this study, UltraSEQ’s sample-level taxonomy predictions (metagenomics module described below) were used for sample-level taxonomy calls, and UltraSEQ’s region-based function and threat prediction were not used. These calculations are provided in the **Supplemental Material B** section for completeness sake as they are useful for other use cases.

#### Metagenomics Service

This service provides sample level taxonomic composition based on the regions identified from reads processed in the query mapper service in 3 steps: 1) filtering out low quality reads, 2) scoring the remaining reads based on the information content of the reads, and 3) predicting the taxonomic composition based on the scores. Sequence region-level taxonomy predictions are associated with confidence scores that are based on alignment quality. For each unique taxonomy identified in the sample, the confidence score from alignments from all query regions that are associated with it are assigned. A K-means clustering approach was used to identify taxonomies with high sample-level scores independently by taxonomy domain (Bacteria, Archaea, Eukaryota, Viruses). Detailed calculations are provided in **Supplemental Material A**.

#### Rules Engine Service

This service combines all of the above context and prediction services for regions, sequences, and samples using user defined logic rules for rapid sequence triage. UltraSEQ currently has 4 default rules engines (Figure 1), but only the MGx Diagnostics Rules Engine was leveraged for this study. The Metagenomics (MGx) Diagnostics Rules Engine enabled flexibility for diagnosis of different infectious disease types (e.g., respiratory, encephalitis/ meningitis, etc.) as outlined in **Supplemental Material E**.

#### Reporting Services

UltraSEQ provides several reports as described below.

#### Top Alignment Report

The top alignment report is a simple report that provides the top alignments from each database (Uniref100 and SoC database for this study) for each query sequence from the Aligner and Query Mapper Services. This report is used for downstream reports as described below.

#### Taxonomy Report

The taxonomy report provides the alignments, annotated with the reference accession’s TaxID, used by the metagenomics service and is used to determine the taxonomic composition of the sample. This report is used for downstream reports as described below.

#### Default Report

The default report provides the UltraSEQ’s regional level predictions for each sequence, including predictions for coarse threat functions (described above), taxonomy, gene ontology, and the sequence-level results of a rules engine(s) if applicable.

#### Sample Report

**Supplemental Material C: File Sample-report_user guide** provides a detailed description of the sample report. The ‘main report’ tab provides a list of all organisms identified from the above Metagenomics Service, the results associated with the identified organisms (relative abundance, number of SoCs identified, etc.), and the metadata associated with the organism from Battelle’s SoC database (whether or not the organism is a human pathogen, whether or not the organism can cause meningitis or respiratory disease, types of respiratory disease, the likelihood that the organism causes disease, whether or not the organism is a common biological or environmental contaminant, references to substantiate metadata, and other information). These results and metadata are used in the MGx Diagnostic Rules Engine (described above) for each disease type studied here (meningitis/encephalitis and respiratory disease), which are summarized in the ‘trigger summary’ tabs. Virulence and AMR factors (both sample-wide and agent-specific) are described in downstream tabs as detailed in **Supplemental Material A**.

### Datasets used for analysis

Several datasets were used during this study as described below.

#### In-silico dataset

Complete reference genomes of 21 different organisms across the tree of life were downloaded from NCBI’s RefSeq database using the open-source bioinformatic tool ncbi-genome-download v0.3.1 (https://github.com/kblin/ncbi-genome-download) with commands as described in their user manual. The downloaded genomes were combined and a few sequences from the Human genome were added to it for testing the host removal process. The combined genomes were used to computationally generate a metagenomic sample using the open-source tool InSilicoSeq v1.5.4 [18] based on the Illumina HiSeq error model.

#### Mock Microbial Community Datasets

To further test UltraSEQ taxonomic profiling capability, we analyzed datasets derived from mixed microbial communities as reported by Nicholls et al. [19]. Specifically, ERR3152364 (nanopore dataset) and ERR2984773 (Illumina dataset) were run.

#### Clinical Datasets

Clinical datasets are summarized in **Table 1**. The datasets represent a range of disease types, sample types, and sequence types. The final datasets (saliva respiratory datasets) were produced as part of this study as described below.

**Table 1.** Clinical Datasets Analyzed in this Study.

| BioProject or Data Source (Reference) | Disease Type | Sample types | Dataset Type | Clinical Data Provided by Study to Compare to MGx Results | Pipeline compared to UltraSEQ |
| --- | --- | --- | --- | --- | --- |
| https://veb.lumc.nl/CliniMG (Xavier et al. [20]) | Encephalitis & Respiratory | CSF, Brain, NP, BAL, Nasal wash, Plasma | DNA and RNA reads | qPCR Positive patient samples (n=13) | 9 Pipelines in Xavier et al [10] |
| PRJNA516289 (Miller et al. [31]) | Encephalitis / Meningitis | CSF | Illumina RNA and DNA reads | Original Clinical Test (n=216 tests) from 95 patient samples** | SURPI |
| PRJNA516582 (Saha et al [32]) | Encephalitis / Meningitis | CSF | Illumina RNA reads | Clinical culture, antigen, or PCR positive specimens (n=36) from patients with neurologic infections based negative specimens (n=30) from patients with alternate diagnosis; CHIKV PCR positive specimens (n=17); confirmed cases from idiopathic set (n=11)*** | ID-Seq with logistic regression model |
| 'CSF_metagenomics' from idseq.net (Hasan et al. [33]) | Encephalitis / Meningitis | CSF | Illumina DNA reads | Culture and/or PCR positive residual specimens (n=74) from patients with suspected CNS infections**** | ID-Seq |
| PRJEB7888 (Fischer et al. [34]) | Respiratory | BAL, sputum, nasal swab | Illumina RNA reads | Influenza PCR positive respiratory specimens (n=24) from patients with seasonal influenza infection and samples (n=5) from patients with pneumonia that tested negative for influenza | Fischer et al. pipeline and Explify |
| PRJNA554856 (Watts et al. [35]) | Respiratory | endotracheal exudate | Ion Torrent DNA reads | Culture positive patient samples (n=2) | Centrifuge and Resistance Gene Identifier |
| PRJNA554461 (Yang et al [36]) | Respiratory | endotracheal exudate | MinION DNA reads | Samples from patients (n=14) with VAP: culture-positive (n=9), culture-negative (n=5), and controls (n=8, patients without pneumonia) | WIMP |
| PRJEB13360 (Graf, Flygare [26, 37]) | Respiratory | Pediatric NP/OP | Illumina RNA reads | PCR-positive samples (n=24); no negative controls | Explify |
| PRJNA634356 (Babiker et al [38]) | Respiratory | NP | Illumina RNA reads | RT-PCR-negative (n = 30) and RT-PCR SARS-CoV-2 and other PCR positive patients with co-infections (n=45) | Explify and KrakenUniq |
| Battelle (this study) | Respiratory | Saliva | Illumina RNA reads | RT-PCR (n=8) SARS-CoV-2 positive patients | N/A |

\* Note: For this case study, 89 samples were run for comparison to Figure 1B in [31]. RNAseq datasets were run for RNA viruses and DNaseq were run for the rest. The 20 “challenge” samples (Supplemental Table 3 in [31]) were not included since the sample IDs could not be matched to raw data files.
\*\* See “All Organisms” in Supplemental Table 2 in [31] (“MNA” IDs match up to raw data sample IDs).
\*\*\* Note: An additional 24 idiopathic meningitis cases (n=24) were reported by the authors [32], but only those with confirmed tests were included in this study.
\*\*\*\* Note: Reads could only be downloaded with host reads subtracted in fastq format (raw files not available). Further, only 20 samples were available. Since most of these are true negatives, 5 samples with confirmed truth tests were selected as a proof-of-concept comparison to Idseq.
CSF = cerebral spinal fluid; BAL, bronchial lavage; VAP = ventilator associated pneumonia; NP/OP = nasopharyngeal, oropharyngeal, N/A = not applicable

### Processing of Saliva Samples for Battelle Dataset

Approximately 1-5 mL specimens were self-collected in RNAase-free 50mL Falcon tubes. Briefly, subjects were instructed to swallow a couple of times prior to collection of normal saliva that naturally pools into the mouth without coughing or sniffing. After collection, the sample tubes were screwed shut and sterilized with a disinfecting alcohol wipe. Samples were submitted to a collection site within 30 minutes of collection where they were immediately placed on ice. Samples were shipped to the laboratory and stored at 2-8°C until diagnostic testing could be conducted. All samples were tested within 72 hours of collection and leftover saliva was stored at −70°C or below for subjects who enrolled in the Battelle Biorepository.

### SARS-CoV-2 qPCR for Battelle Samples

Self-collected saliva samples were analyzed the same day by SalivaDirect™ DualPlex RT-qPCR targeting the N1 gene of SARS-CoV-2 and human RNase P. Briefly, samples were vortexed until homogenous and then 50 µL of saliva was mixed with 2.5 µL of MagMAX™ Viral/Pathogen Proteinase K (ThermoFisher) and vortexed at room temperature at 3000-5000 RPM for 1 minute. Proteinase K was then heat inactivated for 5 minutes at 95°C prior to RT-PCR. The RT-PCR was conducted using TaqPath™ 1-Step RT-qPCR Master Mix according to the SalivaDirect protocol on 5 µL of extraction-free saliva lysate. For each 96-well sample plate tested, a negative template control (NTC) and two positive controls (Twist synthetic SARS-CoV-2 RNA controls at 100 copies/µL) were assessed. Test results were only valid when the NTC returned a negative result, and the positive controls returned the expected positive results for SARS-CoV-2 N1 and negative for the RNase P gene. Detection criteria for clinical samples was as follows: SARS-CoV-2 N1 gene was a C_T_< 37 and the RNase P gene at any value is a positive reportable result. If the RNase P gene had a C_T_<35 and SARS-CoV-2 N1 gene C_T_ ≥40 was a reportable negative result. If the RNase P gene had a ≥35 and SARS-CoV-2 N1 gene C_T_ ≥40 the test was considered invalid and the test repeated for that sample.

### Metatranscriptome Sequencing for Battelle Samples

Saliva from each enrolled subject that returned a positive SARS-CoV-2 qPCR result underwent RNA extraction using the QIAamp Viral RNA (Qiagen) kit for QIAcube according to the manufacturer’s instructions. RNA extracts were then treated using the Kapa Hyper Prep with RiboErase according to the manufacturer’s instructions. Library quantification was conducted by KAPA Library Quant Kit (Illumina) Universal qPCR Mix Kit (Cat# KK4824), the libraries were normalized to 4 nM, and pooled for sequencing on the Illumina NextSeq 500/550 High Output Kit v2.5 (300 Cycles, 150 × 150 bp). FASTQ data was processed through UltraSEQ as described above.

### Positive Predictive Agreement, Negative Predictive Agreement, and Accuracy

Agreement between the metagenomic results with pipeline was compared to reported molecular results (culture, qPCR, antigen testing, etc.) in terms of positive predictive agreement [PPA = TP/(TP+FN)], negative predictive agreement [NPA = TN/(TN+FP)], and accuracy [ACC = (TP +TN)/(TP+FN+FP+FN)]. In cases where negative samples were not available (e.g., Xavier dataset [20]), the additional metric of positive predictive value (PPV) was used for comparisons, where [PPV = TP/(TP+FP)]. Specifically, TPs are defined as instances where the bioinformatic pipeline identifies the same pathogen species as reported by the molecular test; FNs are defined as instances where the molecular test identifies a pathogen but the bioinformatic pipeline does not; FPs are defined as instances where the bioinformatic pipeline identified a pathogen that was tested for, but not identified by the molecular test; and TNs are defined as instances where the bioinformatic pipeline does not identify a pathogen and the molecular test also does not identify the pathogen. If no truth molecular result was available, then the bioinformatic pipeline result was not scored.

For each published dataset and corresponding pipeline, positive and negative calls were documented as reported in the publications, and the detailed results are described in **Supplemental Material D: Supplemental File Scores**. In addition to the pipelines described in the studies in Table 1, for some respiratory datasets, results were compared to Explify results using the Basespace app [21]. For UltraSEQ, positive calls were made if an organism was identified in bin 1A, 1B, 2, or 3 according the logic rules table described in **Supplemental Material E** with the following exceptions. For all datasets, pathogens that rarely cause disease as annotated in Battelle’s SoC database were reported in the raw results but were not considered positive detections. For PRJNA554461, PRJNA516289, PRJNA516582, additional filters were applied due to an obvious *E. coli* contamination. Specifically, for PRJNA554461 *E. coli* was only considered a positive detection if it was identified with >5% relative abundance and >7 SoCs were identified; for PRJNA516582, >50% relative abundance and >21 SoCs filters were applied; for PRJNA516289, *E. coli* positives were not reported under any condition.

## RESULTS

The overall scheme of the UltraSEQ Pipeline providing end-to-end solution for high-accuracy clinical metagenomics is presented in **Figure 1**. Reads are first optionally pre-processed to ensure high quality data. Each read is then aligned to multiple reference databases, and the query mapper server identifies regions in each read. Context services assign taxonomy and functional annotations to each region of each read. The metagenomics module combines information from all reads into a sample-level taxonomy profile. Results are subsequently reported using UltraSEQ’s cloud-deployed web application.

To comprehensively evaluate UltraSEQ’s ability to accurately taxonomically profile metagenomic samples with the end goal of dataset-agnostic diagnostic application, we applied a 3-step evaluation approach. In step one, we evaluated the specificity and sensitivity of UltraSEQ using an *in silico* synthesized metagenome consisting of genomic DNA from 22 different microbial species, including bacteria, fungi, virus, and human. In step two, we evaluated UltraSEQ against a mixed microbial community dataset to measure real world performance. Finally, in step 3, we evaluated UltraSEQ using publicly available clinical metagenomic datasets to compare our results with clinical microbiology test results as well as those from other metagenomic pipelines. The results for each of these comparisons is presented below.

### *In-Silico* Datasets

The information-theory-based approach combined with machine learning algorithms in UltraSEQ allows for a highly tunable final report – ranging from very conservative reporting where false positives are minimized, to more permissive settings which allow for more false positives while ensuring minimal false negatives. Using our *in-silico* synthesized metagenomic sample, we extensively tested the different parameters and found that tuning of UltraSEQ’s Metagenomics Service module (metagenomic module) had the most consequential effect on taxonomic assignment reporting. Within the metagenomic module, this tuning is achieved by altering the user-settable K-means clustering centroid distance tolerance threshold, referred to hereafter as the metagenomic clustering threshold (MCT, **Appendix A**). Our results show that a minimum MCT of 0.5 results in lowest FN counts for the dataset tested, with a very marginal increase in the FP count (**Table 2**). Using a MCT of 0.5, the 4 ‘false positives’ species detected, all at very low relative abundance, were *Aspergillus fumigatus, Lactobacillus paracasei, Prevotella intermedia,* and *Streptococcus equinus*, which indicate reads being assigned from the true positive species *A. niger*, *L. casei, P. denticola* and *S. mutans*, respectively.

**Table 2.** Effect of the metagenomic clustering threshold parameter on specificity & sensitivity of UltraSeq’s taxonomic assignment reporting for our in-silico synthesized sample.

| MCT | True Positives | False Positives | False Negatives |
| --- | --- | --- | --- |
| 0.1 | 18 | 0 | *Human herpesvirus 1, L. monocytogenes, Zika virus* |
| 0.2 | 19 | 2 | *Human herpesvirus 1, Zika virus* |
| 0.3 | 19 | 2 | *Human herpesvirus 1, Zika virus* |
| 0.4 | 19 | 2 | *Human herpesvirus 1, Zika virus* |
| **0.5** | **21** | **4** | **None\*** |
| 0.75 | 21 | 4 | None* |
| 0.8 | 21 | 4 | None* |
| 0.9 | 21 | 4 | None* |
| 1.0 | 21 | 4 | None* |

To compare the taxonomic classification results obtained using UltraSEQ’s metagenomic module with the current gold standard in k-mer mapping-based taxonomy classification tools, we processed the synthetic metagenome sample with Kraken2 [22]. Bar plots comparing taxonomy results for UltraSEQ (MCT 0.5) and Kraken2 are presented in **Figure 2**. While the relative abundance for the False Positive samples were low (< 1%) for both methods, Kraken2 had a higher number of False Positives (6), and also failed to detect *A. niger* in the sample.

**Figure 2.**
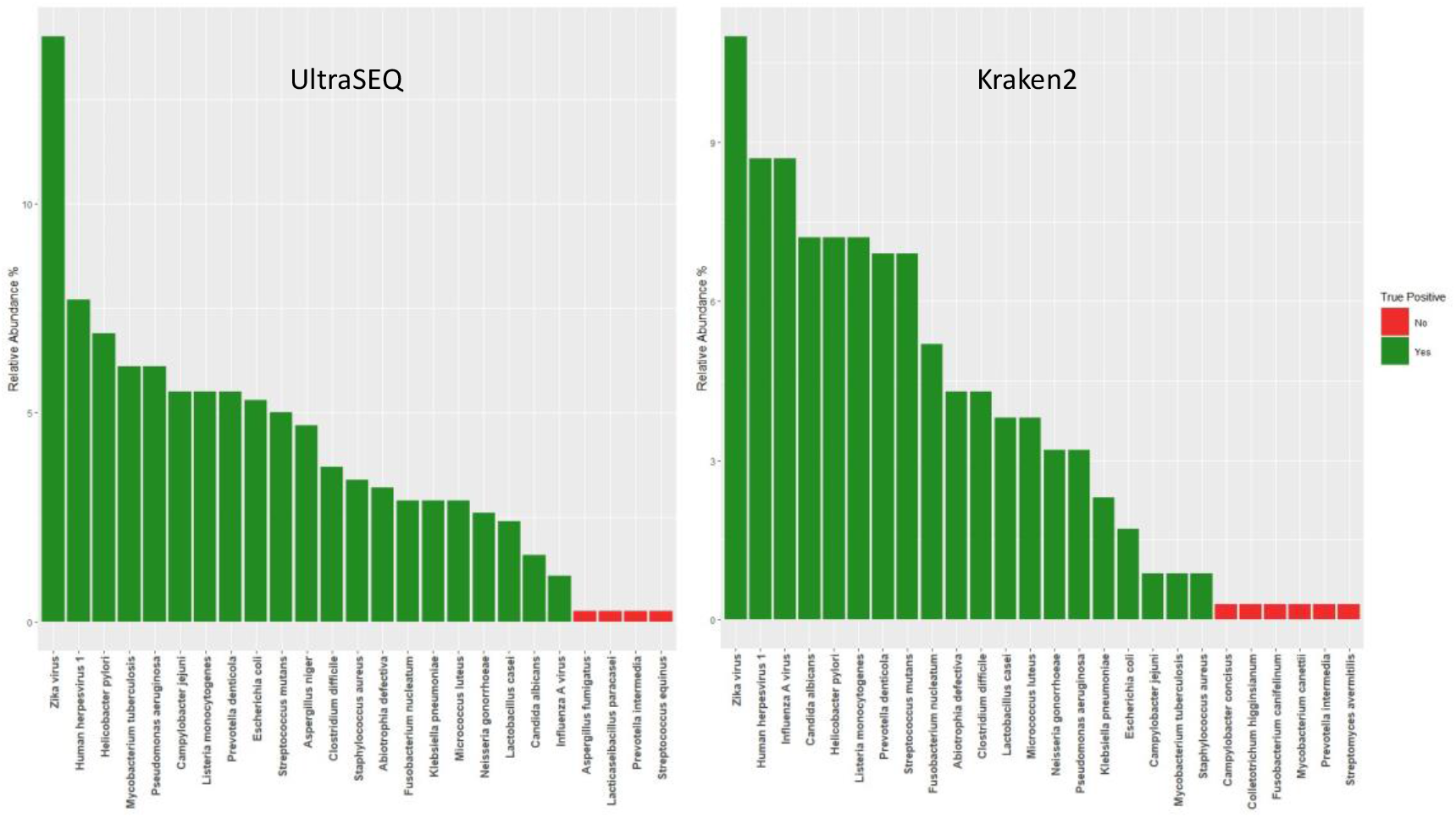
Bar plots showing relative abundance of each species detected from the synthetic metagenome sample using UltraSEQ (left) and Kraken2 (right).

### Mixed Microbial Community Datasets

To further test the metagenomics service with real biological sequences, a mixed microbial community sample, sequenced using both the nanopore and Illumina platforms as described in the methods section, was processed using UltraSEQ. Based on our findings with the *in-silico* dataset, these samples were also processed with an MCT of 0.5. UltraSEQ correctly predicted the presence of all 10 species for both the long and short read datasets with relative abundance values close to expected (**Table 3**). The highest ‘false positive’ species detected in both the datasets was *Bacillus spizizenii*, very closely related to the expected *B. subtilis*. For both datasets, any false positive species detected were much lower in relative abundance (<0.4%) compared to the expected species. These results, and those observed for the synthetic metagenome, led us to apply a domain-specific abundance filter to UltraSEQ’s results for the subsequent application in clinical metagenomics.

**Table 3.** UltraSEQ Correctly Identifies All Organisms in a Mock Microbial Community Dataset

| Species | Expected Relative Abundance % | Illumina dataset: Observed Relative Abundance % | Nanopore Dataset: Observed Rel. abundance % |
| --- | --- | --- | --- |
| Bacillus subtilis | 12 | 11.9 | 8.9 |
| Enterococcus faecalis | 12 | 12.0 | 9.6 |
| Escherichia coli | 12 | 12.4 | 10.5 |
| Lactobacillus fermentum | 12 | 9.0 | 8.9 |
| Listeria monocytogenes | 12 | 12.9 | 14.6 |
| Pseudomonas aeruginosa | 12 | 14.7 | 8.5 |
| Salmonella enterica | 12 | 12.8 | 11.6 |
| Staphylococcus aureus | 12 | 10.4 | 18.8 |
| Cryptococcus neoformans | 2 | 0.5 | 1.5 |
| Saccharomyces cerevisiae | 2 | 1.8 | 2.1 |

### Clinical Datasets

In total, ten different sets of metagenomic datasets, encompassing 407 clinical comparisons from 216 samples were analyzed. These datasets spanned a range of clinical sample types (CSF, nasal, oral, etc.), disease types (respiratory and encephalitis/meningitis), sequence types (RNA and DNA), and sequence generators (Illumina, IonTorrent, and Nanopore). For each dataset, we evaluated UltraSEQ’s diagnostic capability by comparing results from metagenomic datasets of clinical samples to microbiological results from the same samples. For this analysis, we calculated the positive percent agreement (PPA), negative percent agreement (NPA), and accuracy of UltraSEQ with the publicly available sequence datasets or in-house datasets. Overall, UltraSEQ demonstrated an 86% PPA, 97% NPA, and 91% accuracy across all 10 sets of data, demonstrating UltraSEQ’s utility across the wide range of datasets tested. For each dataset, we further compared the performance of UltraSEQ relative to other informatic tools with the aforementioned metrics, as well as additional metrics (antibiotic resistance profiles, usability, etc.) as detailed in the following sections. In contrast to other pipelines, no background datasets were used to generate UltraSEQ results, which is the most direct comparison to clinical microbiological results. All results are shown in **Supplement D**, with each study in a separate tab.

**Figure *3*** shows a summary of UltraSEQ’s clinical diagnostics accuracy against other bioinformatic tools noted for each case study comparison.

**Figure 3.**
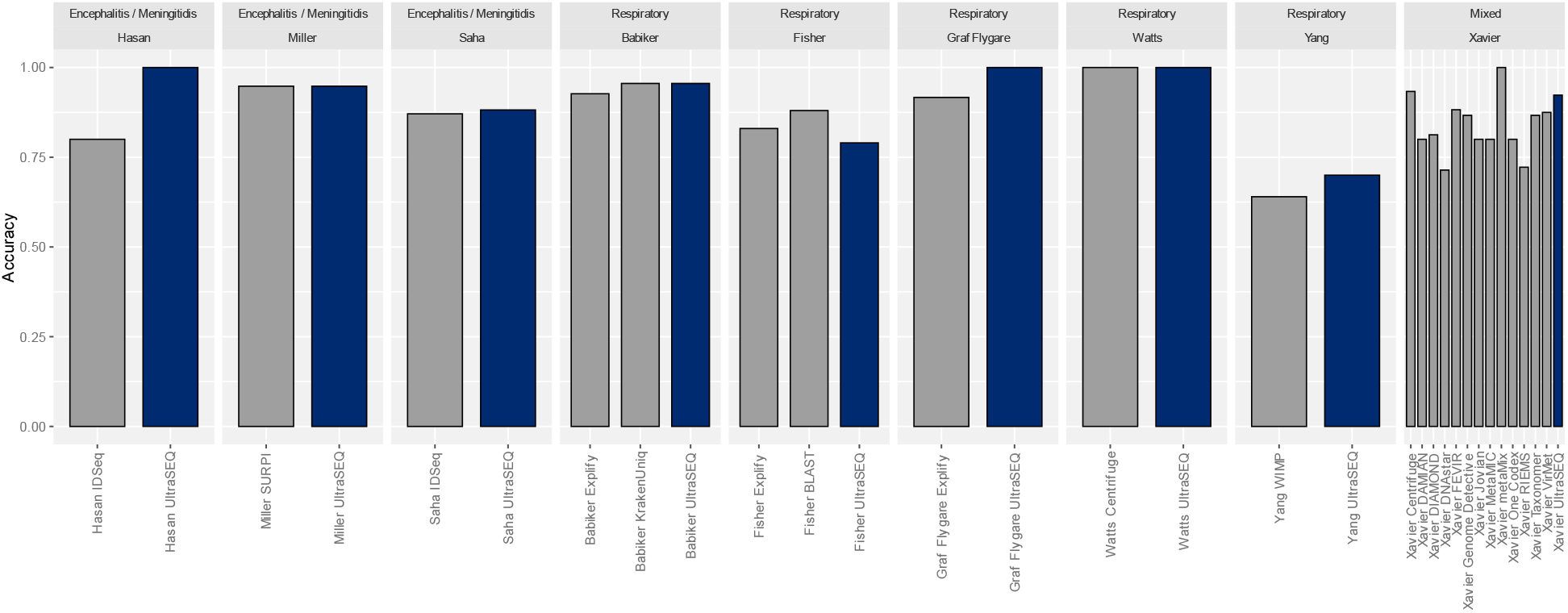
Comparison of UltraSEQ clinical diagnostics performance.

### Case Study #1. PRJNA516289 (Miller et al.)

This study investigated the use of both RNAseq-based and DNAseq-based metagenomics to diagnose encephalitis/meningitis, and the SURPI pipeline was used to make organism calls. For this study, the 90 datasets derived from 76 samples without high host background as reported by the authors were run for comparison. The results for these 90 datasets were compared to 173 clinical and/or confirmatory tests (positive or negative). Overall, UltraSEQ performed the same in terms of accuracy (95%) compared to Miller et al.’s pipeline across all 173 clinical tests (**Table F1**). UltraSEQ performed better than Miller et al.’s pipeline in terms of NPA (100% for UltraSEQ), but not for PPA (85% for UltraSEQ compared to 86% for Miller et. al.). Importantly, UltraSEQ reported zero false positives even without the use of a background subtraction method. Across the various categories tested, UltraSEQ performed better for fungi and bacteria, worse for DNA viruses and RNA viruses, and the same for parasites (**Table F1**). For fungi, Miller et al.’s one false negative was due to lack of identification of *Sporothrix schenckii*, which UltraSEQ identified as a pathogen that can causes encephalitis / meningitis [23]. For bacteria, Miller identified Bacillus species as a false positive in one sample, whereas UltraSEQ did not identify any Bacillus species. For DNA viruses, two false negatives were due to reported co-infections of two different types of herpes virus. For RNA viruses, the two false negatives reported by UltraSEQ that were not reported by Miller, were due to only 2 reads or less being identified as the truth virus, which did not enable high enough confidence in UltraSEQ reporting a positive

### Case Study #2. PRJNA516582 (Saha et al.)

This study also investigated the use of metagenomic RNA sequencing for diagnosis of encephalitis/meningitis. Overall, UltraSEQ performed slightly better than the authors here for the PPA and ACC metrics, with one less false negative (**Table F2**). Further, if the criteria for *S. pneumonia* were loosened from the default criteria described in the Methods Section to only requiring 0.3% relative abundance, an additional 3 FNs would become TPs (88% PPA for all samples). UltraSEQ reported no false positives for all samples (100% NPA), but a direct comparison of NPA values to the authors could not be performed since the authors did not report false positives.

### Case Study #3. ‘CSF_metagenomics’ from idseq.net (Hasan et al.)

This encephalitis/meningitis dataset was selected to compare UltraSEQ to CZ ID (formerly IDseq) [24]. Since most of the 20 available samples were true negatives, 5 samples with confirmed truth tests were selected as a proof-of-concept comparison. For the 5 samples analyzed, UltraSEQ successfully called all 5 correctly (100% accuracy), whereas Hasan called 4/5 correctly (80% accuracy). In the one case where Hasan called a false positive, UltraSEQ did not identify the pathogen (*Streptococcus parasanguinis*). To further demonstrate differences between UltraSEQ and CZ ID, results from analysis of the sample CW322 for UltraSEQ **(Table F3**) and CZ ID (**Figures G1 and G2**) were compared. UltraSEQ only identified two species and automatically filtered out other species with low confidence without requiring user input. In contrast, CZ ID identified many bacterial and viral genera (**Figure G1**), and within each genus, several species were identified. For example, **Figure G2** shows partial results for Neisseria in which 22 species were identified.

### Case Study #4. PRJEB7888 (Fisher et al.)

In this study, Illumina RNA-Seq reads from 24 different respiratory disease patients were evaluated in comparison to the qPCR influenza clinical data. UltraSEQ results were compared to the author’s in-house Blast pipeline and Explify. For this comparison, a positive detection was defined as identification of influenza virus and negative detection was considered lack of influenza virus detection. All pipelines demonstrated 100% NPA, but varied in PPA, with UltraSEQ failing to identify influenza virus in one more and two more cases relative to Fisher et al.’s and Explify’s pipeline, respectively (**Table F4 and Supplemental Material D**).

### Case Study #5. PRJNA554856 *(Watts et al.)*

This case study included samples from two patients with ventilator associated pneumonia (VAP), collected on Day 1 and Day 3, for a total of 4 samples sequenced using the Thermo Fisher Scientific’s Ion Torrent platform. For patient 1, UltraSEQ identified *Staphylococcus aureus* only on Day 3, which is in good agreement with the culture results and author’s results (**Supplemental Material D**). For patient 2, UltraSEQ identified *S. aureus* and *Klebsiella aerogenes* on Day 1, which is in good agreement with the culture results and the author’s results. Other positives were identified as well, including *Prevotella melaninogenica*, *Pseudomonas aeruginosa*, and human alpha herpes virus for patient 1.

#### UltraSEQ for Antibiotic Resistance Profiling

In addition to pathogen identification, UltraSEQ provides antibiotic resistance (AbR) profiles based on the presence of genes that are known to cause resistance to various antibiotics as described in the Methods Section. Results of the AbR profiling for *S. aureus* from sample accession SRR9693434 is shown in **Table F5**. Consistent with the microbial culture results and the author’s results using ResFinder [25]. UltraSEQ identified a profile consistent with resistance of *S. aureus* to methicillin, through detection of the *mecA* gene. The mecA gene was not detected in any other samples, consistent with the authors report. Other potential resistance genes were identified as well, but no other phenotype testing was performed, and thus, no comparative conclusions can be drawn.

### Case Study #6. *PRJNA554461 (Yang et al.)*

This study investigated the use of clinical metagenomics using an Oxford Nanopore MinION sequencer for patients with VAP. Here UltraSEQ results were compared to Nanopore’s What’s in My Pot (WIMP) workflow, as reported in the Yang publication. As shown in **Table F6**, UltraSEQ’s results demonstrated a much better NPA due to 4 less false positives, but a slightly lower PPA due to one more false negative. For the 4 culture negative pneumonia cases (Cases 10 and 12-14) and the 8 control cases (Case 15-22), UltraSEQ results were consistent with the author’s (WIMP) results, with some exceptions as detailed in **Supplement D - Supplemental_File_Scores.xlsx**. For the 10 culture positive pneumonia cases (Case 1-9 and 11), UltraSEQ identified at least one concordant pathogen in 8/10 cases (discordant for Case 7 and 9); in comparison, the authors identified the concordant pathogen in 9/10 cases. However, the authors identified 2 false positives in both Case 1 and Case 4, which UltraSEQ correctly did not report, since its rules engines leverage the metadata associated with each identified sample.

#### Antibiotic Resistance Profiling

To further test UltraSEQ’s ability to identify AbR profiles based on genotypes, UltraSEQ results were compared to the phenotype results reported in Yang et al, who used ResFinder [25] (**Table F7).** In general, UltraSEQ showed excellent agreement with phenotypic results, and the results provide a direct interpretation of genotypes (i.e., UltraSEQ automatically interprets antibiotic resistance based on CARD hits, whereas Yang et al. required manual interpretation based on hits to various genes). Specifically, for 4 Cases, antibiotic resistant phenotypes to 11 antibiotics were identified using culture profiling; of these 11 antibiotics, UltraSEQ identified *pathogen-specific* evidence (i.e., the resistance genes likely derived from the identified pathogen) for resistance for 7 of those antibiotics or classes of antibiotics. For 3 of the remaining 4 (all fluoroquinolone antibiotics), UltraSEQ identified *pathogen-agnostic* evidence (i.e., the gene was not identified to be associated with the identified pathogen); for the remaining 1 of 4, UltraSEQ identified *pathogen-specific* evidence to a closely related antibiotic (different type of beta lactam antibiotic).

### Case Study #7. PRJEB13360 (Graf, Flygare)

These studies investigated the use of Illumina RNA sequencing of 24 upper respiratory tract samples to diagnosis diseases caused by respiratory viruses. No negative controls were included in this study, and all results were compared to PCR tests. For these 24 samples, Flygare et al. reported an average of 95% sensitivity (positive predictive agreement) for their Protonomer module of the Taxonomer software (average as reported in Figure 3 in [26]). In contrast, UltraSEQ identified the correct virus in all 24 samples (100% PPA). Since the individual viruses that UltraSEQ identified in each of the 24 samples were not reported by Flygare et al., we performed a head-to-head comparison for a subset of the samples of UltraSEQ to Explify, the software developed from the Taxonomer software [21]. For the 12 samples analyzed, Explify correctly identified 11/12 (92% PPA), whereas UltraSEQ identified 12/12 (100% PPA) (see **Supplemental Material D**). For accession ERR1360082, Explify failed to identify Enterovirus B. The results for all other samples were very similar.

### Case Study #8. PRJNA634356 (Babiker et al [9])

This study investigated Illumina RNA sequencing datasets from nasopharyngeal (NP) swabs for the diagnosis of SARS-CoV-2 infection. Pathogen detections were compared to SARS-CoV-2 RT-PCR-positive patients (n=45) and RT-PCR-negative subjects (n = 30), including one viral co-infection confirmed by RT-PCR. We compared UltraSEQ’s results to the author’s, who used KrakenUniq [27], and a subset were also run using Explify. Molecular and confirmation testing used by the authors detected viruses such as influenza, respiratory syncytial virus (RSV), parainfluenza, human metapneumovirus (hMPV) and rhinovirus, and/or human coronaviruses (HCoV). Thus, a positive detection by any pipeline was only scored if it detected one the above viruses and the specific test was run for that sample. As compiled in **Table F8 and in Supplement D,** UltraSEQ’s results demonstrated a better PPA compared to both pipelines due to no false negatives. In contrast, Babiker reported one FN (GA-EHC-084F) and Explify software resulted in 3 FNs. For FPs, UltraSEQ identified two in contrast to one FP reported by Babiker et al. and zero by Explify. Taken together, the results demonstrate that UltraSEQ has an overall accuracy of 96% for this dataset compared to 96% and 93% for Babiker’s pipeline and Explify, respectively.

### Case Study #9. (Xavier et al.)

This study compared 13 different bioinformatic pipelines for the diagnosis of respiratory and encephalitis from metagenomic datasets. For this dataset, UltraSEQ showed a 100% PPV (no false positives) and 92% PPA (sensitivity), as shown in Table F9. This PPA value was better than 10 of the other 13 pipelines, but only 1 (metaMix) of the 3 pipelines that had better PPA values had an equivalent PPV. NPV or accuracy was not calculated here since no true negative samples were included in the dataset.

### Case Study #10: In-house Covid-19 Dataset (PRJNA856680)

To further test UltraSEQ’s ability to diagnose respiratory disease with a different sample type (saliva), we generated and analyzed 8 RNAseq metagenomic datasets from COVID-19 positive patients. These results were compared to gold standard PCR tests (**Supplement D**). For 6 of the 8 samples, we correctly identified SARS-CoV-2 (75% true positive rate). No other pathogens (except for oral/throat bacterial contaminants) were identified. For the two false negatives, no reads were identified that aligned to SARS-CoV-2, and the lack of detection did not correlate with the qPCR results, suggesting the lack of detection was due to lack of signal in the dataset and not due to UltraSEQ.

## DISCUSSION

Applied metagenomics is a powerful emerging capability enabling untargeted detection and characterization of pathogens, reducing the time window for, and enhancing the identification of, emerging and traditional threats for both clinical and biosurveillance applications. Realizing the potential for metagenomics requires bioinformatic solutions that can not only keep pace with scientific discovery but are also scalable, accurate, and science-backed. While clinical metagenomic tools have been developed and validated for specific applications, their utility is limited by workflow-specific requirements, computational constraints, lengthy expert review, and stagnant databases. For example, SURPI has been validated for CSF samples/ encephalitis using Illumina reads [1], Explify for respiratory samples/ disease using Illumina reads [4,7,8], and Karius for cell-free DNA/ sepsis from Illumina reads [28], but each of these requires specific wet laboratory workflows, instrumentation, and background controls. In parallel, numerous metagenomic tools have been developed for taxonomic identification with potential utility for surveillance such as CZ ID [24] and What’s In My Pot (WIMP) [29], but such tools will likely not have widespread clinical application until validated across various use cases.

Here we presented results for evaluation of our novel UltraSEQ pipeline using *in silico* datasets, mock microbial community datasets and publicly available clinical datasets across a wide range of applications. For the *in silico* dataset, UltraSEQ successfully detected all 21 species across the tree of life with fewer false positives than Kraken2 (Figure 2). UltraSEQ also detected all 10 bacterial and fungal species in the mock microbial community dataset sequenced via both the Illumina and Nanopore platforms (Table 3). Finally, the clinical datasets contained samples from different infection types (encephalitis, meningitis, other respiratory diseases), comprised of both short-read (Illumina, IonTorrent) and long-read (Nanopore) sequencing data in both RNA-Seq and DNA-Seq formats, and represented both sterile (e.g., spinal fluid, blood) and “dirty” (e.g., saliva, nasal) sample types. Here, UltraSEQ’s pathogen detection accuracy was the same or better than the comparable bioinformatic tools used in the studies for seven of the nine clinical datasets (**Figure *3***). UltraSEQ analyzed the set of diverse samples and identified pathogens and achieved an overall accuracy of 91%, even without requiring dataset-specific configuration settings changes, background sample subtraction, or prior sample information. Further, we demonstrated UltraSEQ’s ability to provide accurate antibiotic resistance and virulence factor genotypes that are consistent with phenotypic results.

Taken together, the above results demonstrate that Battelle’s UltraSEQ platform offers a transformative approach to microbial and metagenomic sample characterization, employing biologically informed detection logic, deep metadata, and a flexible system architecture for classification and characterization of taxonomic origin, gene function, and user-defined functions, including disease-causing infection. A highly curated pathogen and virulence factor database underpins the UltraSEQ analytics engine and enables rapid, accurate, and explainable detection and characterization of pathogens. In addition to the results shown here, we further discuss the important features of the UltraSEQ software compared to both the benchmarked software described in the results section as well as additional well-known software used for clinical diagnostics and/or surveillance research (**Figure 4**). While each of the twelve pipelines provides advantages and disadvantages, UltraSEQ’s curated databases, logic-based approach, and modularity provide accurate predictions across a wide range of dataset types with best-in-class comprehensiveness of reference database coverage, science-backed annotations, and flexibility.

**Figure 4.**
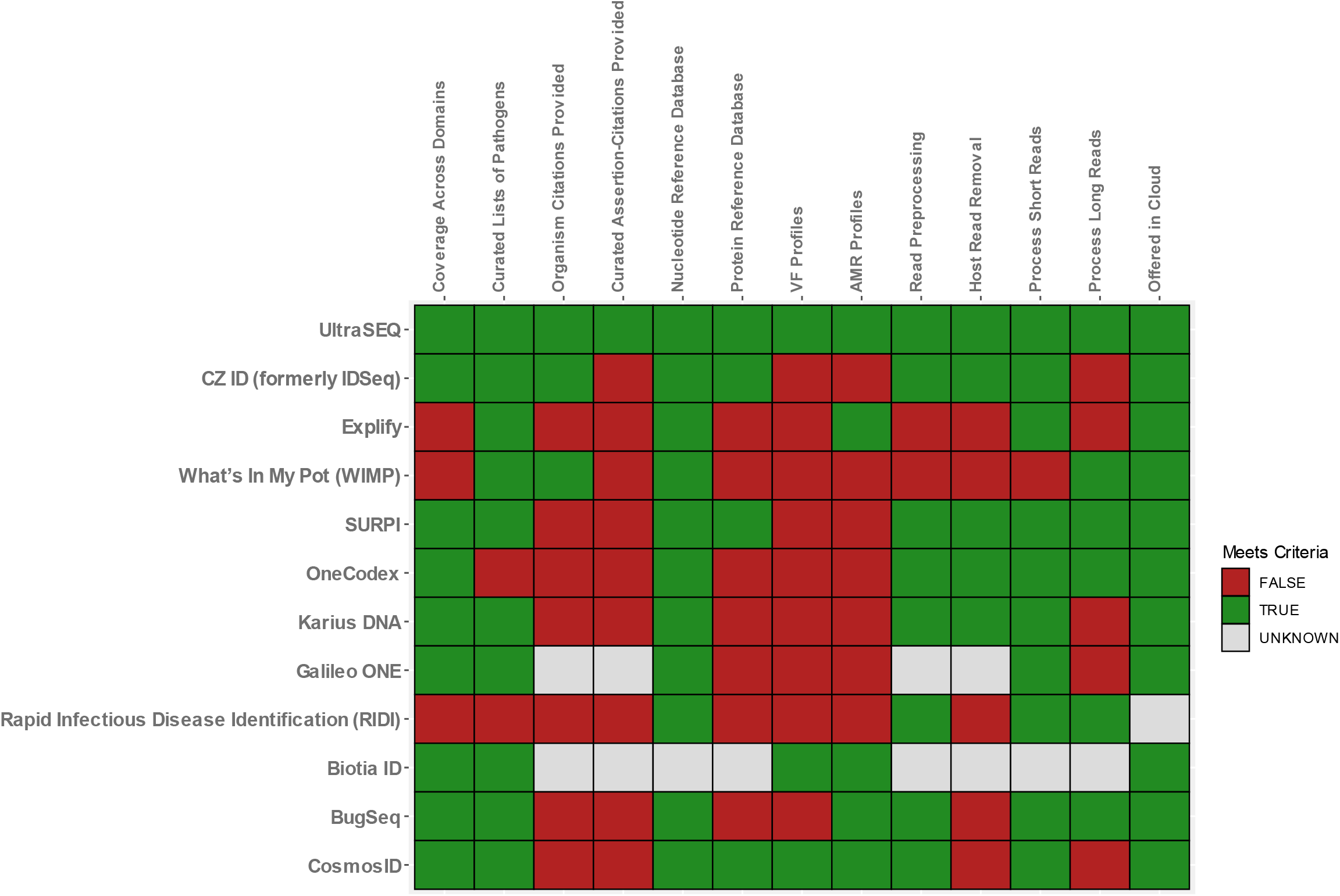
Comparison of UltraSEQ to commonly used bioinformatic tools for clinical diagnostics or surveillance [21, 24, 29, 30, 39–50].

### Comprehensiveness

Metagenomic-based identification of pathogens relies on matching query sequences to sequences in a database, followed by prediction. To ensure the highest accuracy, maximal coverage across the tree of life is required to ensure that query reads are not incorrectly assigned. For example, non-typical parasites that can cause disease, such as *Trichinella* spp. (identified in the Miller et al. dataset) can only be identified if they are included in subject databases. When comparing the twelve tools, UltraSEQ is one of the nine tools to have taxonomy coverage across all relevant biological domains (bacteria, eukaryote, virus, and fungi). Of the three tools that do not have complete taxonomy coverage, Explify and WIMP are missing eukaryote and Rapid Infectious Disease Identification (RIDI) is missing virus, fungi, and eukaryote domains. Further, UltraSEQ is one of only four tools that allow both nucleotide and protein reference databases to be included in analysis pipelines, expanding the quantity of reference sequences that can be used to classify samples. In addition to taxonomic coverage, UltraSEQ’s databases include coverage and provide both virulence factor and antibiotic resistance genes like two other tools (**Figure 4**).

### Science-back Annotations

While database coverage is important for pipeline performance, curation of the data contained in the database is perhaps even more important. All but two tools leverage curated lists of pathogenic organisms to enable specific reporting of pathogens identified in a sample. For example, SURPI is backed by annotated lists of pathogens that cause encephalitis/meningitis, Karius is backed by lists of sepsis-causing pathogens, and Explify is backed by lists of respiratory disease causing pathogens, including granularity on phenotypic groups (normal flora, colonizers, etc.). UltraSEQ provides these curated lists as well, but provides additional granularity, including likelihood of the pathogen to cause disease (e.g., immunocompromised), contaminant type (e.g., from laboratory reagents versus from native flora), specific disease subtypes (e.g., ventilator associated pneumonia), etc.

In addition, providing end users with human-readable text and primary references relating to organisms, gene functions, and other information can minimize the burdens (such as review time) associated with bioinformatic predictions. Based on our research, UltraSEQ and at least two additional tools provide such text and citations. For example, CZ ID provides background information on organisms and provides Wikipedia citations, and WIMP provides background information on organisms at the genus level [30]. However, to our knowledge, UltraSEQ is the only tool that provides curated assertion-citations of pathogens. Specifically, we provide end users with not only a short text rationale (assertion) for why the data is curated in a particular way, but also the primary reference (e.g., PubMed ID) in order to provide highly explainable and accurate results. For example, in the Yang et al. dataset, WIMP identified *S. maltophilia, Staphylococcus epidermidis, Pseudomonas aeruginosa,* and *Klebsiella pneumoniae* for Case 1, but only *S. maltophilia* was the likely disease-causing organism based on culture results. UltraSEQ identified *S. maltophilia* and *S. epidermidis*, but *S. epidermidis* could be eliminated because of its metadata (lack of association with VAP and known skin contaminant) and the reference for this annotation is provided to the end user.

### Flexibility and Accuracy Across Datasets

Like some other tools, UltraSEQ provides the functionality to both preprocess and remove host-derived reads. While most other tools perform read preprocessing, at least five do not include a host read removal step, which can increase computation time. Further, UltraSEQ and four other tools can process both short (e.g., Illumina, IonTorrent) and long (e.g., Nanopore, PacBio) read sequencing data. Further, UltraSEQ provides the additional flexibility of being deployed in different formats, including cloud deployment.

Most importantly, unlike the implementation of other tools shown in this paper across the clinical case studies discussed, we have demonstrated that implementation of UltraSEQ for clinical metagenomics for different sample types does not rely on parameters that are specific to **read depth** and **background subtraction**. For example, in their implementation of CZ ID, Saha et al. used specific filters that were dependent on read depth (e.g., NT reads≥10, NR reads ≥ 2, etc.). For background subtraction, Miller et al. used a 10X threshold above the background sample [31] with their implementation of SURPI. Similarly, with their implementation of CZ ID, Hasan et al. and Saha et al. used (among other filters) an empirically derived Z score based on test sets of samples [31, 32]. In contrast, UltraSEQ uses an information theory approach to calculate positive detections utilizing the metagenomic module results and its own rules engine. Thus, the predictions are not dependent on read depth and no background sample subtraction is required to achieve a high degree of specificity, allowing UltraSEQ to produce excellent results for any type of dataset without *a priori* knowledge.

To further illustrate UltraSEQ’s lack of reliance on arbitrary depth thresholds and a background sample, we ran several datasets at various read depths and saw no degradation (or improvement) in UltraSEQ’s performance (data not shown). Further, we noticed that *E. coli* was a pervasive contaminant in many of these datasets, including the Hasan et al. and Miller et al. datasets. For these datasets, UltraSEQ also identified *E. coli* in most samples. Unlike the authors who used several custom user-defined filters and background samples to remove this contaminant, UltraSEQ was able to leverage its metadata-aware logic engine to easily identify *E.coli* as a false positive with great confidence, and thereafter automatically eliminate it from the result without requiring background data or additional human intervention. Overall, UltraSEQ made all of its predictions without a background sample, maintaining comparable or better accuracy to the studies that utilized a background sample subtraction analysis step. Like qPCR and other molecular assays, we suggest that background samples should be included as separate controls (and not used as a background subtraction). Therefore, we believe this feature of UltraSEQ will be a major benefit in its clinical metagenomics application since UltraSEQ does not require specific empirical testing to remove background signals and apply specific thresholds, thus expanding utility across any sample type.

Future development of UltraSEQ will focus on improving performance classifying samples co-infected with multiple pathogens in the same domain (e.g., two viruses). For example, in the Fisher et al. dataset, UltraSEQ identified a false negative (ERR690513). For this sample, human alphaherpes virus dominated the relative viral abundance maps (see Methods section); thus, UltraSEQ likely did not report influenza virus in the case due to a low number of influenza virus reads relative to human alphaherpes virus reads. Another area of future development would be towards expanding and optimizing UltraSEQ’s ability to detect and classify emerging novel pathogens. Recent work in our laboratory analyzing novel SARS-CoV-2 demonstrated that UltraSEQ could identity SARS-CoV-2 as a SARS-CoV-1 related virus (data not shown). Further, analytical models built from hazardous function signals (using the methodology reported in our recent publication [9] demonstrated that SARS-CoV-2 clusters along with other SARS-CoV viruses with similar host range and human receptor type. Future work will expand upon these advances and improve UltraSEQ’s ability to characterize features of clinical and research significance from emerging pathogens. Finally, Battelle is committed to continued curation of the pathogen databases at the heart of the UltraSEQ to ensure relevance and advantage over contemporary tools. We believe Battelle’s UltraSEQ will emerge as the leading platform for clinicians looking to leverage the promise of modern metagenomics in solving the unrelenting challenges of infectious diseases diagnosis.

## Conflict of Interest/Disclosures

### Data and Software Availability

Metagenomic reads the saliva samples from this study have been submitted to the NCBI BioProject database (https://www.ncbi.nlm.nih.gov/bioproject) under accession number PRJNA856680. The UltraSEQ platform is available for use at https://www.ultraseq.org/, with accounts available through contacting the authors.

### Supplementary Materials

This manuscript contains the following supplemental materials:

Supplement A – Detailed UltraSEQ Services

Supplement B – Other UltraSEQ Services

Supplement C – Sample-report_user guide

Supplement D – Supplemental_File_Scores

Supplement E – UltraSEQ Rules Engine Logic

Supplement F – Supplemental Results

### Author’s Contributions

BG, CM and CB conceived the idea for this manuscript and wrote the majority of the of the manuscript. <u>BG and CM performed the bioinformatic analysis and generated the visualizations.</u>

PF implemented UltraSEQ on AWS and locally and processed all datasets. CH curated the majority of the data used in processing. RS, LC, AS, and AMS contributed to all laboratory work, including sample extraction, sequencing, qPCR analysis, and methodology reporting for SARS-CoV-2 saliva samples. <u>All authors read the final manuscript and have approved.</u> BG and CM performed the bioinformatic analysis and generated the visualizations.

## Acknowledgements

A special thanks to Brett Fowle, Omar Tabbaa, Nathan Parry, and Robert Wieczorek for their development efforts to the UltraSEQ software and curation applications. This work was supported in part by the Centers for Disease Control and Prevention, via contract number 75D30121C11398, as well as Battelle internal research and development funding.

